# Defensiveness measurement in honey bees (*Apis mellifera*) and brain expression of associated genes after noxious stimulus

**DOI:** 10.1101/2022.04.15.488528

**Authors:** Jenny P. Acevedo-Gonzalez, Alberto Galindo-Cardona, Nicolas L. Fuenzalida-Uribe, Alfredo Ghezzi, Tugrul Giray

**Affiliations:** Department of Biology, University of Puerto Rico; CCT NOA SUR, National Scientific and Technical Research Council (CONICET), Argentina; Miguel Lillo Foundation, Tucumán, Argentina

**Keywords:** Africanized honeybees, Gene expression, gAHB

## Abstract

Honey bee (*Apis mellifera* sp.) colonies and individuals respond variably to disturbances. In this study, we examined the role of neural modulation and metabolism in constitutive and experience-dependent differences in defensive response. We compared brain gene expression in bees of identified gentle and defensive colonies in a standard assay. For neuromodulation, we examined membrane receptor genes for the biogenic amines dopamine (DOPR2), octopamine (OAR), and serotonin (5HT2a), and the enzyme gene in the synthesis pathway (THR). To examine neural metabolism, we assessed the Oxidative Phosphorylation Pathway “OXPHOS” gene expression (i.e., ND51 and ND20-LIKE). Bees of defensive colonies had a significantly lower expression of amine receptor, synthesis genes, and OXPHOS genes. Experience differences or exposure to nociceptive neurons activated by nocive stimuli (electric shock) led to differences in the expression of all genes except 5HT2a. The same target genes demonstrated an increase in expression levels after electric shock and sting response. We discuss the convergence of neuromodulation, neural metabolism

## Introduction

Honey bees, as other organisms, can gate defensive behaviors to defend vital resources such as brood, food, shelter and reproduction, which are fundamental for their survival (Alcock, 2009). Uniquely, the *Apis mellifera* L., as a model animal, exhibits great advantages for behavioral studies (Seeley 1996, Galizia et al. 2012) of social organization. The phenotype of the functional colony depends on both the collective of individuals and the characteristics of each individual (Page et al., 2012, Avalos et al., 2020). The Africanized gentle bees (gAHB) found in Puerto Rico (Rivera Marchand et al., 2012) are a good study model for understanding how honey bees respond to perturbations at the level of colonies and of individuals.

In the highly structured societies of bees, the individuals constantly modify their behavior in response to the social signals of their congeners (Traniello and Robinson 2021). For instance, only a subgroup of individuals shows aggressive defense of the colony, that is stinging and pursue of intruders (Rittschof, 2017). The differences in the worker’s behavior in response to a stimulus at the colony level is thought to rely on genetically and developmentally explained differential response thresholds to particular stimuli by different workers (Robinson, 1987 and 1992; Duarte et al., 2012; Jeanson and Weidenmüller, 2014). Changes related to age, genetics, and adaptation to the environment have been studied as mechanisms underlying honey bee defensive behavior (Robinson and Page, 1988; Kolmes and Fergusson-Kolmes, 1989; Giray et al., 2000; Hunt, 2007; Alaux et al., 2009). The regulation of the expression of defensive behavior relies on the environment in which the bees live, as well as on their physiological, genetic, historical, and evolutionary characteristics (Bloch and Grozinger, 2011). In colonies, only a subgroup of worker bees of 14 days or older takes on guarding tasks. Guards examine other bees or insects entering the colony and intercept possible intruders. Likewise, a subgroup of workers with 2-3 weeks of age (called guardians) shows aggressive defense and may sting vertebrate enemies (Nouvian et al., 2016). The colony-level differences in defensiveness are thought to have a greater portion of individual workers performing these defensive tasks (Avalos et al., 2014). In addition, bees may respond defensively based on previous experience, both at the individual and colony levels (Avalos et al., 2014). Thus, bees in a colony that experiences an episode of aggressive defense will remain ready to respond for up to three days, and their brain gene expression profile is different from bees not engaged in aggressive defense (Alaux et al., 2009). Mechanisms underlying variation in readiness to defense may differ for constitutive differences and experience-dependent differences (rev. in Traniello and Robinson 2021). At the individual level, there are several common neurophysiological mechanisms related to aggression in vertebrates and invertebrates. In most vertebrate animals, in addition to neuropeptides such as vasopressin (Ferris et al., 1997) and steroid hormones such as testosterone and cortisol (Fox et al., 1997), the neuromodulation by biogenic amines is important for defensiveness (Evans, 1986; Veroude et al., 2016). These molecules are neuromodulators of the nervous system of vertebrates as well as invertebrates (Evans, 1986; Monastirioti, 1999; Blenau and Baumann, 2001), where Dopamine (DA), Noradrenaline (NA), and Serotonin (5HT) are linked to regulation of defensive behavior (Summers and Greenberg, 1995; Huber et al., 1997; Korzan and Summers, 2021; Pandolfi et al., 2021). Octopamine (OA), described as a functional analog of NA in invertebrates, is involved in invertebrate defensive behavior (Roeder et al., 2003).

Dopamine (DA), serotonin (5HT), and octopamine (OA) levels measured in the brain of honey bees are associated with age, age-related division of labor, and task specialization (reviewed by Giray et al., 2007). Older bees working outside the hive present higher values of amines than the bees that work within the hive, storing food and caring for the offspring (Schulz and Robinson, 1999). Older bees take on high-risk tasks such as foraging and defense (see Alaux et al., 2009).

Other mechanisms are demonstrated to be important, at the least correlated with defensive behavior in bees (Alaux et al., 2009). Oxidative Phosphorylation pathway “OXPHOS” genes in the bee brain are implicated in differential gene expression studies. OXPHOS genes differ across “killer” AHB (African Honey Bee) versus docile EHB (European Honey Bees) (Alaux et al., 2009).

We hypothesize that differences in this biogenic amine and OXPHOS genes underlie defensiveness differences also within a population of Africanized honey bees in Puerto Rico. The regulation of the expression of defensive behavior relies on the environment in which the bees live, as well as their physiological, genetic, historical, and evolutionary characteristics. Since this is part of the running hypothesis as to why the bees in PR are different. We selected six candidate genes based on previous work and gene expression patterns of bee races (Alaux et al. 2009). We predicted constitutive differences in these genes would be correlated with the defensiveness of honey bee colonies. In addition, we predicted manipulation in the form of application of harmful stimuli would lead to regulatory changes in these same genes. We first examined the correlation of gene expression levels of colonies classified as highly defensive and gentle within the gAHB population. Later we examined whether the defensiveness-related genes change in response to harmful stimuli, i.e., electric shock that induces Sting Extension Response SER (Sandoz et al. 2007, Agarwal et al. 2011).

## Materials and Methods

### Collection site

Samples of gAHB workers were collected from hives maintained at the Experimental Apiary at the Gurabo Agricultural Research Station of the University of Puerto Rico (18.25732, - 65.9866).

Bee population in Puerto Rico has a uniform genetic distribution and the Experimental Apiary houses ca. 40 hives that are representative of this population (Galindo-Cardona et al. 2013).

### Experiment 1. Comparison of Gene Expression Levels in Bees From Colonies with Differences in Defensiveness

First, we evaluated the defensive behavior of each hive following the protocol used by (Giray et al., 2000; Guzmán-Novoa et al., 2003), and more recently described in (Avalos et al., 2014; Avalos et al. 2020). Each behavioral test was done twice with at least four days between tests; the values (see Avalos et al., 2014) obtained for each hive were averaged. Each test began by applying two puffs of smoke (a way for calming bees) to the entrance and one additional puff of smoke to the top once the outer cover was removed to open the colony. One frame was lifted, and the intensity of the corresponding behaviors was recorded according to four categories: run, sting, hang, and fly. The lowest intensity for each behavior is given a score of 1, and the highest a 4. In this test, the minimum score possible is 4, and the maximum is 16. We determined the score for each hive, and for this study, we selected the two most defensive (Defensiveness Rank= 12) and the two least defensive colonies (Defensiveness Rank = 4). After identifying the defensiveness scores, colonies were left undisturbed for 1 week. From ranked colonies we collected a total of 20 individual worker honey bees from each of the two most and the two least defensive colonies and immediately placed them in liquid nitrogen, and transported them to the laboratory for processing (see below). We analyzed the brain expression levels of OA1, DOPR2, 5HT2a, TRH, ND51, and ND20-like genes for both colonies.

### Experiment 2. Changes in Gene Expression Levels Following a Sting Response

For the second experiment, bees from the two colonies with the highest defensiveness level were analyzed in their gene expression. We test whether, after the electric shock stimulation, the bees receiving electric shocks (treatment group) change the expression of analyzed genes in contrast to control bees (untreated group) that were not exposed to electric shock, only placed in the electric shock box. We analyzed the brain expression levels of OA1, DOPR2, 5HT2a, TRH, ND51, and ND20-like genes both in control and treatment individuals. We collected groups of workers from each of the two previously selected colonies and transported them to the lab. These groups were kept overnight in an incubator at 34°C, 80% RH (Percival incubator, model I-30NL). The following day, we examined the individual response of each worker by using the sting assay paradigm previously described by Avalos et al. (2014). The apparatus consisted of a grill formed by steel electrodes with 3mm spacing (see Figure 1, in Avalos et al., 2014), connected to a power source (BK Precision 0-30V Analog) to provide a shock when the circuit is closed by a bee. The bee remains between the steel grill and a glass cover, held 5mm above the grill with two side rests made of thick Plexiglas. A thin cover of petrolatum jelly is applied to the glass cover to prevent bees from turning and holding on to the glass cover. This way, bees are forced to be in contact with the steel grill continuously. The protocol consists of exposing bees to short (10 seconds) bouts of continuous electric shock at a set voltage with a brief resting period (20 seconds) between each bout. Bouts of shock was provided on an increasing scale from 0 V until the bee attempted to sting the apparatus, with each subsequent bout being 2 Volts higher than the previous one (e.g., 0, 2, 4, etc.). Once a bee attempted to sting components of the grid, it was removed from the setup and immediately frozen in liquid nitrogen. As a control group, we used bees from the same colony that were exposed to the apparatus but received no shock over a 2-minute interval and then frozen in liquid nitrogen. All samples were stored in a −80 °C freezer for further processing.

**Figure 1.**
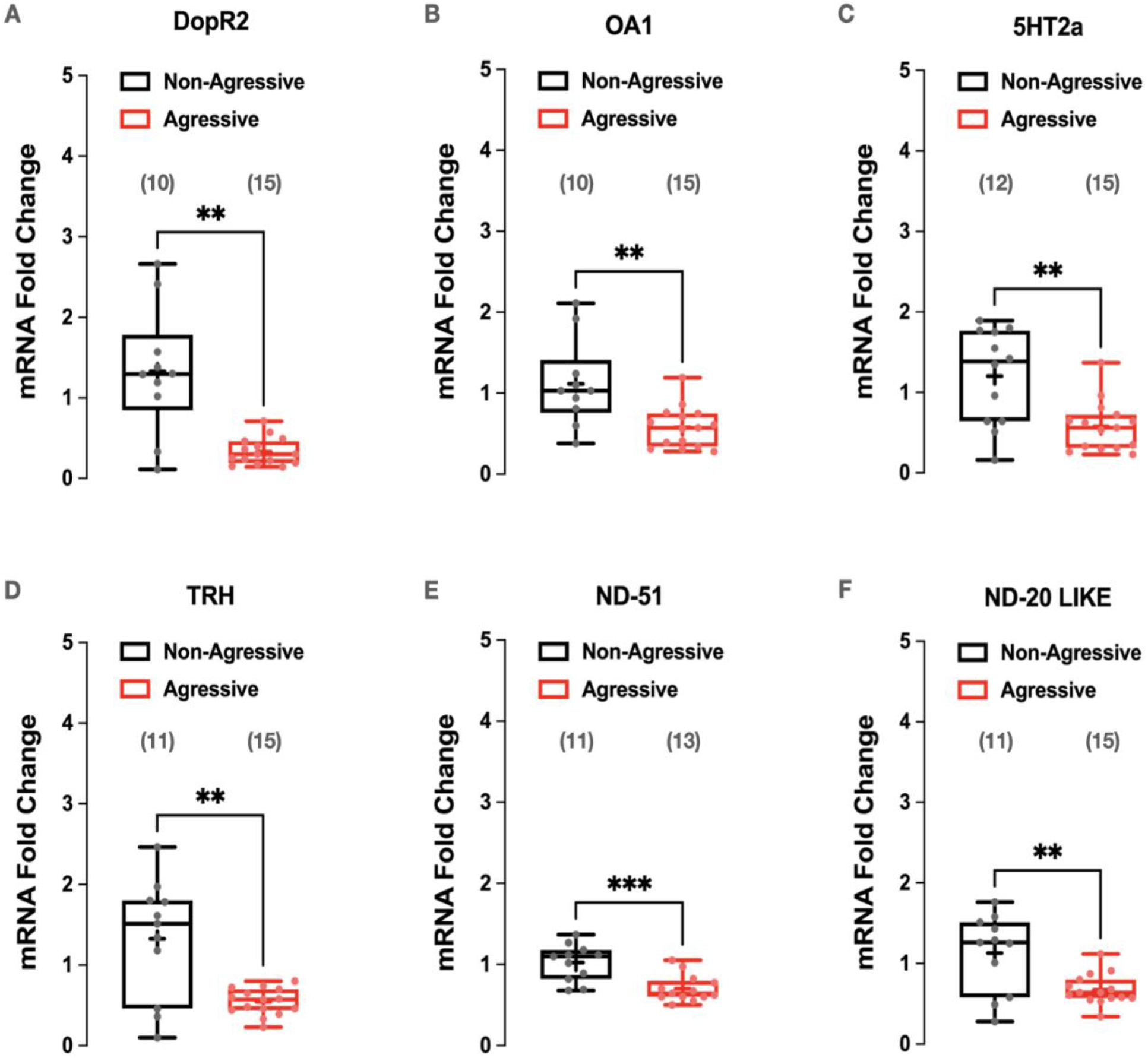
Gene expression levels across docile vs defensive colonies. Gene expression of aminergic receptors (A-C), serotonin biosynthesis pathway enzyme (D) and OXPHOS pathway genes (E-F) into the brain of docile and defensive gAHBs. Gene Expression levels are normalized using Rps5 gene as housekeeping reference. Number of samples are shown between parentheses. Welch’s t-test was performed using GraphPad Prism, * = P<0.0332, ** = P<0.0021, *** = P<0.0002 and **** = P<0.0001.

#### RNA isolation

The sets of bees were frozen in liquid nitrogen and then stored in a −80 °C freezer until the time of dissection. For each bee, heads were removed, and brains were dissected over dry ice (Avalos et al. 2021). The extracted brain was then placed in RNALater^®^-ICE (Invitrogen) until RNA extraction. Brains were placed individually in 1 ml of Trizol and macerated to break down tissue. After the addition of 100 μl of Bromo-3-chloropropane (BCP), samples were briefly vortex and let sit at RT for 15 min, then centrifuged at 21,000 g for 15 min at 4 °C to generate an aqueous phase. The clear upper layer was transferred to a fresh tube on ice. After the addition of (500 μl) of isopropyl alcohol, the aqueous phase was again vortexed and left at RT for 10 min, then centrifuged for 10 min to produce a pellet. The pellet was subsequently washed with (500 μl) of 75 % ethanol prior to drying and resuspended in DEPC water. The pellet was incubated at 65°C for 10 minutes, kept on ice, and then stored at −80 °C. The purity and concentration of RNA were determined by Nanodrop 1000. The quality of the extracted RNA (integrity and size distribution of total RNA) was verified by 1.2% agarose gel electrophoresis in the TAE buffer and gel star staining system.

#### cDNA synthesis

Aliquots of the samples were organized in a 96well PCR plate and reverse transcribed to cDNA using the iScriptTM Reverse Transcription Supermix kit and protocol (BioRad; Hercules, CA) following the manufacturer’s instructions. The program of reverse transcription consisted of a cycle of 5 minutes at 25 °C (hybridization), a cycle of 20 minutes to 46 °C (reverse transcription) and stopped at 4 °C. cDNA was diluted 1:2 with sterile water prior to qPCR analysis.

Gene expression levels were measured using qRT-PCR with a STRATAGENE 7900 detector and the SYBR green detection method (BioRad). A housekeeping gene, rpS5, was used as a loading control. It was used a thermal profile of 95°C for 30 s followed by 95°C for 5 s and 60°C for 30 s. Steps two and three were repeated for a total of 40 cycles and included plate reads for fluorescence during each step at 60°C. Following the cycle program, products were denatured for 10 s at 95°C., re-annealed and then a dissociation profile was measured between 69 and 95°C at an increment of 0.5°C to provide evidence for reaction fidelity (Evans P. 1986). Positive and negative control reactions were run on each 96-well plate.

#### Target Genes

In this study, we used a panel of genes to examine transcriptional signals associated with defensive behavior. The reference gene used was Ribosomal Protein S5 (rpS5) which has been previously validated as a reliable reference gene (Evans P. 1986; Evans J. 2004; Evans and Wheeler, 2001). This target gene panel was composed of six candidate genes. Four of these genes are associated with biogenic amines and two genes associated with the Oxidative Phosphorylation Pathway (OXPHOS) previously identified in studies of colony aggression (Li-Byarlay et al., 2014).

This methodology allows determining the degree of specificity to observe the different patterns of expression of some receptors of Biogenic Amines: Octopamine Receptor (OR1), Dopamine Receptor (DOPR2), Serotonin Receptor (5HT2a), Tryptophan Hydroxylase (TRH) and Oxidative Phosphorylation Pathway: ND20-like and ND51. Three biological and technical replicates were prepared. The test was performed using GraphPad Prism version 7.04 for Windows.

#### Primer Design

Reference gene primers were obtained from previously published sources (Evans P. 1986; Evans J. 2004; Evans and Wheeler 2001). The target genes and primer sets were developed in a multi-step process using Integrated DNA Technologies Primer Quest^®^ and Oligo Analyzer computational tools. Sequences of target homologues were obtained from databases at the National Center for Biotechnology Information (NCBI). For each candidate gene, sequences were input into IDT’s Primer Quest^®^ tool together with specific search parameters detailing primer (17-20 bp) and amplicon (150-200 bp) size, as well chemical characteristics of the amplicon and qRT-PCR reaction (e.g., 58-62 ^0^C product Tm, 3 mM Mg^+^ concentration, etc). A summary of primers used is shown in Table 1.

**Table 1.**
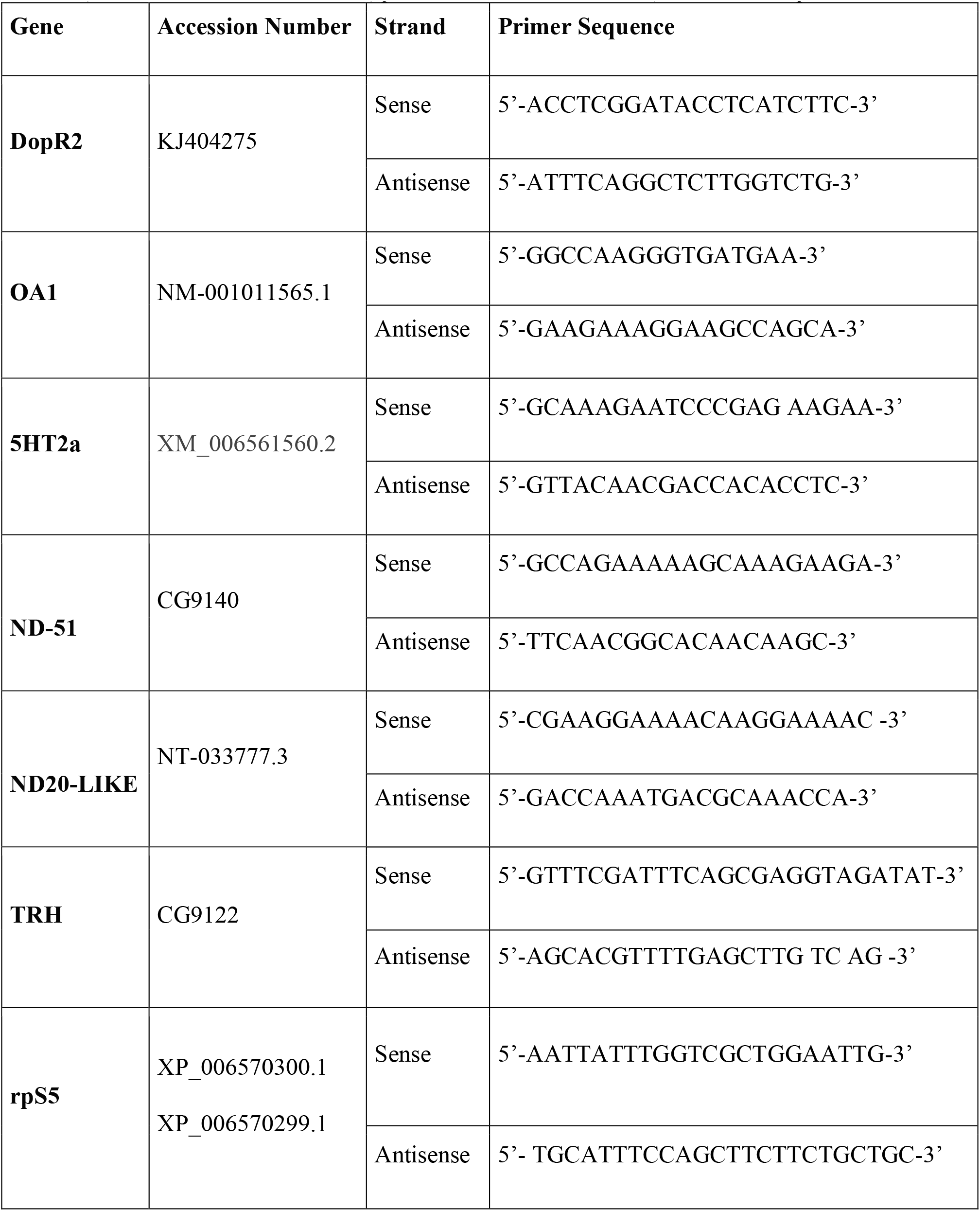
Primer Sets Used for Each of the Target Candidate Genes and Housekeeping. Reports primer sequences for the set of genes used. The information provided are the genes’ name, GenBank accession number, primer strand read direction, and actual sequence.

#### Quantitative Real Time Polymerase Chain Reaction

Optimized primer sets were used together with iTaq™ Universal SYBR^®^ Green Supermix (BioRad) and aliquots of the samples to conduct our qRT-PCR analysis. For each gene, three 96-well PCR plates were run in a Stratagene™ MX3005P qPCR system. Plates were prepared and run on the same day with template, primers (Table 1), and corresponding SYBR Green Supermix at the 10 μL total volume reaction level (SB Green 5 μl, 10 μM Primer F and R and 1 ul of cDNA template) resulting cycle thresholds (C_t_) were checked and samples that did not produce at least two consistent values across the three plates were discarded from the study. Although rare, replicates were also discarded if the product T_m_ did not match expected amplicon T_m_ value, as this is an indication of possible miss-priming or primer artifacts during amplification.

#### Data Analysis

Raw Ct’s were first normalized using the 2^-(Ct(target)-Ct(reference))^ method and the resulting target gene expression level relative to rpS5 expression level was analyzed by using standard ΔΔCt method Livak and Schmittgen, 2001; Nolan et al., 2006). The first analysis compared non-defensive groups (docile) across the brain for each gene to determine if relative target gene expression levels were constant or different. A Student’s t-test was conducted to compare the gene expression associated with a docile behavior type against defensive behavior in the selected bees. The second analysis in sting responsiveness to electric shock of increasing voltage (2, 4, 6, 8 until 20 seg). We compared relative expression levels of the treatment group (sting responses) with the control group (untreated) to determine if expression levels significantly differed between groups.

## Results

### Experiment 1. Comparison of Gene Expression Levels in Bees From Colonies with Differences in Defensiveness

We identified a range of defensiveness levels across colonies in the apiary. The two least defensive (docile) and the two most defensive colonies (defensive) were used for gene expression analyses. For all genes examined, the number of transcripts detected was higher in the brains of docile bees than in the brains of defensive bees. (see Figure 1).

### Experiment 2. Changes in Gene Expression Levels Following a Sting Response

We found that five of six analyzed genes were significantly different between control and treatment bees. The levels of dopamine and octopamine receptors (i.e., DOPR2, OA1) were higher in the group that received shock treatment (Figure 2), thus indicating signaling from both aminergic systems. Similarly, TRH, ND51 and ND20-Like genes also exhibited increased expression levels after electric shock. In contrast, 5HT2a, showed no significant differences in expression levels between groups (Figure 2C), thereby suggesting serotonin is involved in the general awareness of the colony but not in the individual response to a harmful stimulus.

**Figure 2.**
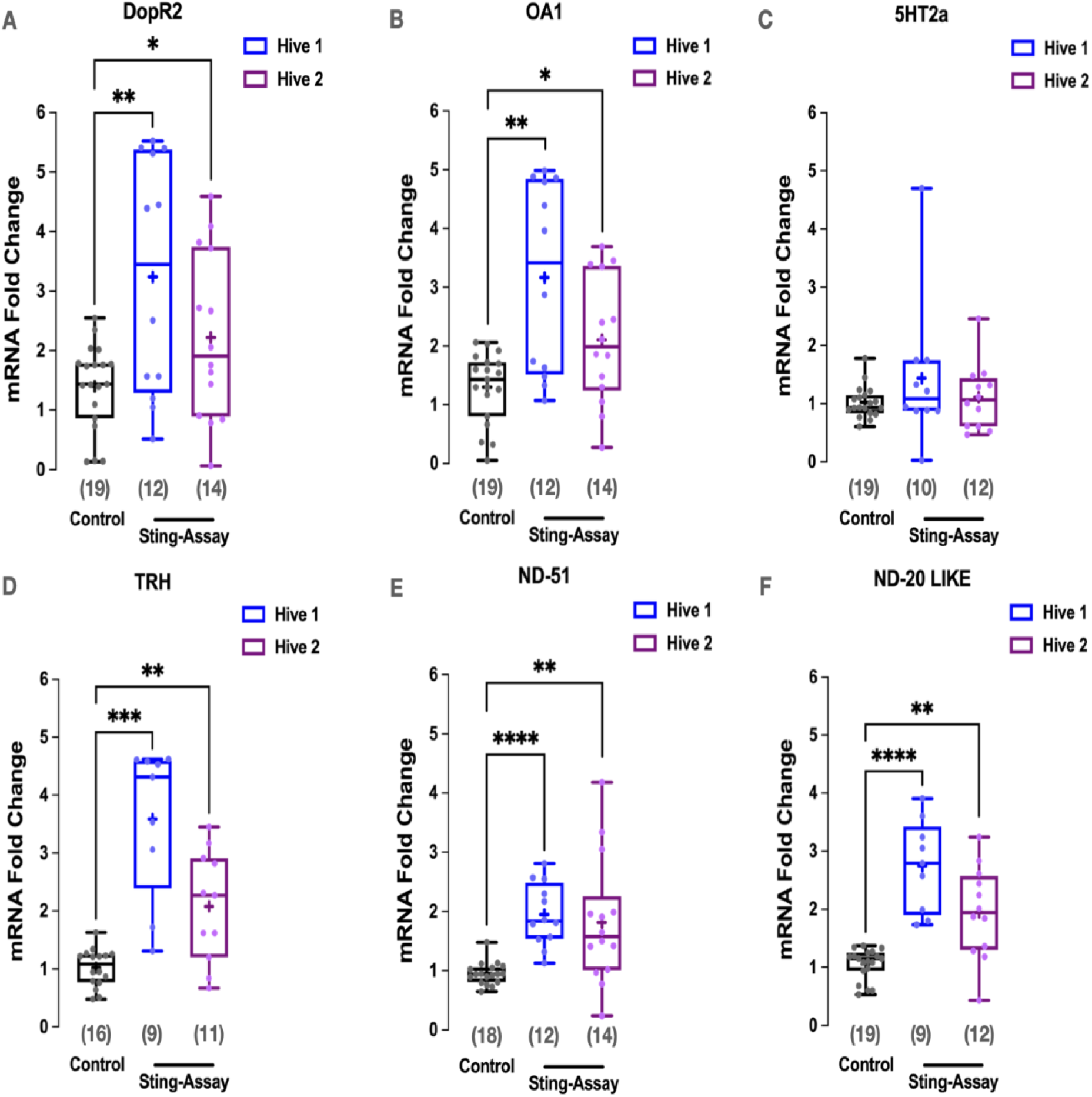
Gene expression levels in response to manipulation or harmful stimuli in two defensive colonies. Expression level of aminergic receptors (A-C), serotonin biosynthesis pathway enzyme (D) and OXPHOS pathway genes (E-F) of two hives (defensive colonies) were subjected to electric shock (Sting-Assay) or not (Control). N=20 per treatment. Number of samples are shown between parentheses. Welch’s t-test was performed using GraphPad Prism, * = P<0.0332, ** = P<0.0021, *** = P<0.0002 and **** = P<0.0001.

## Discussion

The main finding of this study was that the biogenic amine pathway and brain oxidative metabolism are important both for colony differences in defensiveness and individual responses to nocive stimuli. The baseline expression level of all examined genes in these pathways differed across colonies with notable differences in defensive behavior. In addition, expression levels of all but one gene (**5HT2a**) changed in response to a harmful stimulus. Both pathways and multiple genes were shown to be associated with defensive behavior in separate studies previously (Alaux et al. 2009, Chandrasekaran et al. 2015), in this study all examined candidates were found to be related constitutively or facultatively to defensive response. Moreover we compared bees within the same population, known as gAHB (Rivera-Marchand et al. 2012) or Puerto Rico Honey Bee (Feliciano-Cardona et al. 2020). Focus on a single population may have provided the opportunity to examine mechanisms that may be hidden with large contributors to differences when bees of different subspecies or populations such as EHB vs AHB (Breed, Guzmán-Novoa & Hunt, 2004) are contrasted.

### Comparison and Correlation of Baseline Gene Expression Levels and defensive behavior

Even in bees sampled at rest, (and not identified as foragers or soldiers), or before any defensive response we found differences in target genes, consistent with known role of biogenic amines and OXPHOS pathway genes in honey bee defensiveness. The inhibition of oxidative phosphorylation increases defensive behavior and this has been associated with a shift toward aerobic glycolysis (Li-Byarlay et al., 2014). The experimental manipulations described by Li-Byarlay and colleagues (2014) mimic constitutive changes in brain gene expression, similar to reduced oxidative metabolism in brain disease associated with increased aggressiveness. In this study, these constitutive differences in gene expression levels across high and low defensive bees are in agreement with the idea that the colony or social background is important for defensive behavior (Li-Byarlay et al. 2014, Nouvian, Reinhard and Giurfa, 2016, Avalos et al. 2020).

### Sting Response and Change in Gene Expression Levels

Interestingly the facultative changes in expression of Dopamine and octopamine receptors in response to electric shock stimuli are opposite to differences of constitutive level of expression observed between docile and defensive gAHB colonies. This suggests basal lower expression levels for these genes underlie the greater neuromodulation in response to aversive stimuli, with a large range of response. In fact, studies that compared bees that respond to harmful stimuli match the pattern observed in this work, with bees engaged in defense to have higher expression levels of the target genes (Alaux et al. 2009). Similarly, TRH, ND51 and ND20-Like genes also exhibited increased expression levels after electric shock, suggesting these are also involved in this process. In contrast, 5HT2a, showed no significant differences in expression levels between groups (Figure 2C), thus suggesting serotonin is not involved in facultative response directly; yet it may influence the general awareness of the colony, thereby rendering bees ready to respond.

Various hypotheses have been raised on the mechanisms that modulate defensive behaviors in honeybees (Hunt, 2007; Breed, Guzmán-Novoa & Hunt, 2004): e.g., environmental, developmental, context (bee in the colony context or as individual under lab conditions) and genetics (Breed, Guzmán-Novoa & Hunt, 2004). In terms of gene expression, differences have been observed between defensive AHB and docile EHB, across age groups and across presence and absence of defensive response (Alaux et al., 2009). In addition, pharmacological approaches showed biogenic amines modulate the defensiveness in bees (Tedjakumala et al., 2014) and the differences of defensive/aggressive behaviors among bee colonies correlate with gene expression of biogenic amine and OXPHOS pathways (Alaux et al., 2009; Chandrasekaran et al., 2015; Hunt, 2007; Nouvian et al., 2018). Particularly, studies have shown changes into mitochondrial activity after a harmful stimulus (Rittschof et al., 2019). Biogenic amines are demonstrated to modulate responsiveness to nocive stimuli (Nouvian et al., 2018; Rittschof et al., 2019). These separate studies provided the targets examined in the current study, and generally results are consistent with previous studies.

It has been shown that during defensive behaviors oxidative metabolism is increased in the bee’s brain (Rittschof et al., 2019), thus indicating neuronal metabolism via oxidative phosphorylation. This may suggest a shifting metabolic pathway preference in the body versus the brain. In a context of stressful situations, as in the protection of the hive from intruders, bees exhibit short, fast flights to escape or to attack. These efforts, that might as well end in sacrifice of the bee, may redirect the cellular energy to quick release from glycolytic pathway in the body. The defense behaviors such as short, fast flights require increased cellular metabolism (Giray et al., 2000; Chandrasekaran et al., 2015). As a trade-off, the brain may switch to slower energy production through increased OXPHOS metabolism in the brain.

It would be interesting to determine whether there are regulatory mechanisms underlying such a trade-off in metabolism across brain vs body. To our knowledge, no previous study has compared gene expression at the level of the brain with gene expression at the level of the body (e.g., fat body, thorax, among others) during defensive response. We do see the opposing levels of expression of OXPHOS pathway in constitutive vs facultative responses in this study. These observations are in agreement with similar observations and experiments with opposing findings matching constitutive (Li-Byarlay et al. 2015) or facultative contexts (Rittschof, Rubin & Palmer, 2019). If this tradeoff between brain and body occurs, a negative correlation would be predicted for the target gene expression in the brain and the rest of the insect body. Interestingly, recent studies that examine tissue-specific gene expression or epigenetic changes in social insects demonstrate head-thorax-abdomen or brain-muscle-fat body differences in relation to reproduction and diapause or aestivation (Döke et al., 2015; Rasmussen et al., 2016, Breshanan et al. 2022).

## Conclusion

Defensive behaviors comprise various biological factors: including neurobiological, genetic, hormonal, nutritional and brain chemistry processes. However, biological factors alone do not determine expression of defensive behavior. The social environment of the individual is also important in controlling underlying neurobiological processes and behavior (Liu, 2004, Li-Byarlay et al., 2014). Defensive behavior is a consequence of the regulation of external and internal stimuli in the organism (Breed, Guzmán-Novoa & Hunt, 2004). Our findings demonstrate the importance of both biogenic amines signaling and cellular metabolism, in the expression of defensive behavior in gAHB. These could be further explored to understand evolutionary changes in defensive behavior of the Puerto Rico Africanized bee population.

## Acknowledgements

We are indebted to the members of our laboratories as anonymous reviewers by the comments. The UPR-Sequencing-Genotyping Facility contributed for analyses. Humberto Ortiz-Zuazaga and Jose C. Bonilla assisted in the HPCf facility. Thanks to Arian Avalos for his suggestions during the development of the research and Carolina Monmany and Yarira Ortiz for her suggestions in the writing.

